# Mathematical modeling of navigational decisions based on intensity versus directionality in *Drosophila* larval phototaxis

**DOI:** 10.1101/248856

**Authors:** Lucia de Andres-Bragado, Christian Mazza, Walter Senn, Simon G. Sprecher

## Abstract

Organisms use environmental cues for directed navigation. Depending on the sensory modality and complexity of the involved sensory organs, different types of information may be processed. Understanding the basic logic behind navigational decisions critically depends on the complexity of the nervous system. Due to the comparably simple organization of the nervous system of the fruit fly larva, it stands as a powerful model to study decision-making processes that underlie directed navigation. Here, we formulate a stochastic method based on biased Markov chains to model the behavioral basis of negative phototaxis. We have quantitatively measured phototaxis in response to defined sensory inputs. We find that larvae make navigational decisions by taking into account both light intensities and its spatial gradients, and our model allows us to quantify how larvae minimize their exposure to light intensity and at the same time maximize their distance to the source of light. The response to the light field is a non-linear response and saturates above an intensity threshold. Our mathematical model simulates and predicts larval behavioral dynamics only using light intensity and directionality as input parameters. Moreover, it allows us to evaluate the relative importance of these two factors governing visual navigation. The model has been validated with experimental biological data yielding insight into the strategy that larvae use to achieve their goal with respect to the navigational cue of light, paving the way for future work to study the role of the different neuronal components in this mechanism.

**Author Summary:** Navigational decision-making is a complex process during which the nervous system is able to decipher external input through molecular and cellular mechanisms to produce a spatially-coordinated behavioral output. *Drosophila* larvae provide an excellent model to understand these decision-making mechanisms as we can measure the behavioral output (larval navigation) in response to quantifiable external input (different light conditions). We have performed experiments to quantify larval light avoidance in order to subsequently design a mathematical model that quantitatively reproduces larval behavior. Our results allow us to characterize the relative importance of light intensity and directionality and yield insight into the neural algorithms used in the decision-making mechanism of larval phototaxis.

## Introduction

The nervous system is functionally organized to perceive external cues, which are encoded and decoded to make the correct behavioral decisions. A way of studying the logic of these decision-making mechanisms is through the analysis of robust stereotypical navigational strategies evoked by controlled stimuli (such as light or odor cues) in simple model organisms. Indeed, much of our understanding in the fundamental logic of taxis has been achieved by studying organisms such as *Escherichia coli, Caenorhabditis elegans* or larvae of the fruit fly *Drosophila melanogaster* (1, 2). Navigation in *Drosophila* larvae has been assessed in response to different types of single sensory inputs including vision (3, 4), olfaction (5, 6, 7, 8) and thermosensation (9, 10, 11) and a combination of inputs to study multisensory integration (12, 13).

*Drosophila* larvae show robust navigation towards appetitive and away from aversive cues. However, in the absence of an external cue or for a group of blind larvae, the dynamics is essentially random-like (14). In the presence of a light source, we experimentally show that the taxis is still moderately stochastic, being at most three times less efficient than ballistic dynamics. Therefore, an underlying Markov chain presents itself as an excellent basis for modeling larval dynamics since the external field (light) only introduces a small perturbation that allows a quasi-equilibrium description. That Markov chain is biased to take into account the external cue using Boltzmann’s probabilities (15). Measurable magnitudes can be obtained by averaging them over the simulated trajectories. Stochastic techniques like the one we are proposing here have been recently used to model taxis as a diffusion problem (16). The power of such stochastic techniques rests on its capacity to tackle complex problems, like diffusion or phase transitions, with a moderate cost in computational time. In particular, the Metropolis-Hastings chain has been used to efficiently locate global minima of combinatorially-complex objective functions such as the *travelling salesman* problem (17). In this work, we show how to introduce generalized Metropolis-Hastings weights to bias a Markov chain to extract information from biological experiments where larvae take decisions using information gathered from their immediate surroundings.

Previous approaches to model taxis behavior have taken advantage of dividing the animal’s movements into a set of discrete behavioral states and to analyze the transitions between these states. One modeling approach has been based in a linear non-linear Poisson cascade to model the transition between the larval states (12, 13). Another approach has been to model larval taxis as continuous oscillations whose direction is not controlled by the stimuli but by an *intrinsic oscillator* and where the external stimuli would influence the amplitude of the oscillation (18).

*Drosophila* larval phototaxis provides an excellent model to study behavioral decision-making because their exposure to the sensory stimulus of light can be tightly controlled (3, 4). Their paths bear a strong connection with the intensity of light and to the position of the light source, which in the literature has been termed as light directionality (3). Larvae perceive light through a pair of bilateral eyes, which have been shown to be absolutely essential for visually-guided navigation (4, 19, 20, 21, 22, 23). Information from the external field of light is then processed in the brain by a genetically hard-wired decision-making algorithm. Our model allows us to characterize that algorithm as a combination of a goal-directed behavior layer on top of a stochastic one.

By focusing on the impact of two key excitation elements, light intensity and light directionality, we present a model that studies the interplay between these two components for navigation. We exploit experimental navigational data obtained from various controlled illumination conditions to define probability weights that we use to polarize an underlying Markov chain that has been introduced to analyze larval taxis using stochastic methods. Such a mathematical model allows us to simulate larval dynamics with a minimal number of free parameters. Our model provides a theoretical and experimental framework of the decision-making mechanism functioning in the larval brain during navigation.

## Results

### Impact of light intensity and its spatial gradient on taxis

The navigation index (*NI*) has been used in the literature as a significant statistically-averaged dynamical parameter describing sensory guided navigation (3, 24). Along a given direction (*x*), the is defined as the mean velocity in that direction, *v*_*x*_, divided by the total velocity in any direction, *v*. In other words, since velocities are measured over a common time interval, the *NI* in a certain direction can be taken as the distance moved in that direction, Δ*x* divided by the length of the stratified path, *s*. For example, the *NI* along the *x* axis, *NI*_*x*_, would be defined as 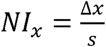 (Fig 1A). Therefore, the value of the *NI* provides an assessment of the efficiency of larval navigation. In our case, since the source of light was located in the +*x* axis of the agarose plate, *NI*_*x*_ the would be approximately –1 for an object moving ballistically away from the illumination source (negative values for the *NI* imply that larvae navigate towards the -*x* axis). On the other hand, we would obtain a *NI*_*x*_ of approximately for a random walk taken in the absence of an external cue or for blind larvae.

**Fig 1.**
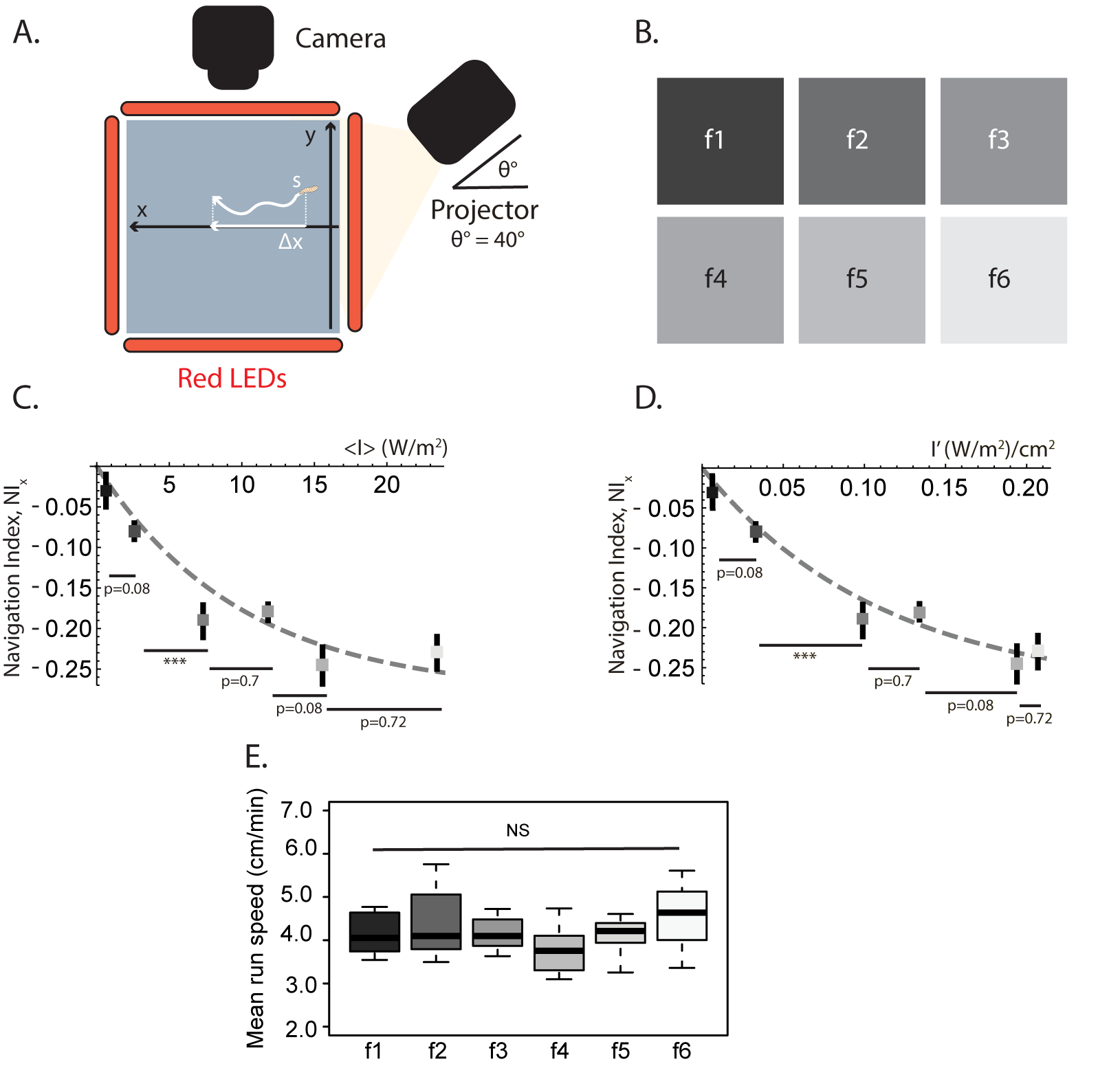
Larval navigation depends on the absolute light intensity and the gradient of the light field. (A) Experimental set up formed by the agarose plate (*x*–*y* plane), light source (projector located at *x* = 36.5, *y* = 0, and *z* = 25), forming an angle *θ* = 40° with respect to the agarose plate, and a videorecording camera. The larvae move on the agarose plate and their positions are recorded by the camera. The LEDs placed on the border of the plate aid in the image-acquisition process. The navigation index in the *x* direction (*NI*_*x*_) is calculated by dividing the distance moved in the *x* axis, Δ*x*, by the total length of the stratified path, *s*. (B) Filters with different light intensities and gradients, f1 – f6 (S1 Fig1 and Table 1). (C) *NI*_*x*_ and standard deviation plotted against the average intensity over the plate for each filter, *I*< (W/m^2^); the resulting curve shows that the efficiency of larval navigation depends on *I*<. The dashed line is a least-squares interpolation to data used to guide the eye (see text). The values for the different filters were compared using the Welch t-test and Benjamini-Hochberg was used to correct for multiple comparison. (D) Same as (C) as a function of the gradient for each filter, *I*′ (W/m^2^/cm^2^). (E) Larval mean run speed for the filters f1-f6 takes values from 4.0 cm/min to 4.5 cm/min, which is a statistically non-significant difference (NS). The whiskers of the boxplots represent the range of the mean run speed for the different filters. Within the boxplots for each filter, the middle band is the median (50^*th*^ percentile), and the length of the boxplots shows the 1^st^ and 3^rd^ quartile (25^th^ and 75^th^ percentile).

Previous studies have shown that the intensity of light, its spatial gradient and light directionality are relevant factors for visually-guided navigation (3). Therefore, we hypothesize that these components may be sufficient to explain the observed biological behavior in our defined experimental framework. First, we experimentally tested the dependence of our primary dynamical measurable magnitude (*NI*) with the two independent variables determined by the external light field: the absolute light intensity and its gradient over the agarose plate where the larvae are located (intensity is measured using irradiance units, as the radiant power flux received per unit area, see Materials and Methods). For this, we used a set of filters where light intensities and their spatial gradients have been varied in a controlled and gradual way: f1, f2, f3, f4, f5 and f6 (Fig 1B). For all these experiments, the angle of the source of light was kept constant at 40°. Our measurements show that navigation of wildtype larvae depends on the absolute light intensity. Larvae show a very low navigation score when the light intensity is low, such as in f1, where *NI*_*x*_ = ‒0.03. As the light intensity increases, the value of the increases in a non-linear way and it saturates for intensities higher than *I* > 20W/m^2^ to a value of around *NI*_*x*_ = –0.3 (Fig 1C). A heuristic expression that interpolates the dependence of the measured with the absolute light intensity can be written as:

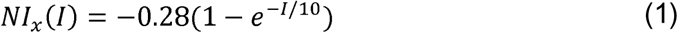

This interpolating function captures the two salient features of this experiment. Firstly, as expected for a dynamic process based on a Markov chain, it approaches zero in the absence of external stimulus (*I* = 0). Secondly, it saturates for around *I* ≈ 20 W/m^2^ (Fig 1C).

Moreover, larval navigation also depends on the slope of the filter, *I*′ (Fig 1D).Same as for the light intensity, in the absence of a gradient, navigation is quite random-like,*NI* (*I*′ ≈ 0) ≈ 0, and it shows saturation for gradients steeper than *I*′ > 0.20 W/m^2^/cm (Fig 1D). A heuristic interpolating function that only depends on the gradient of the field of light is:

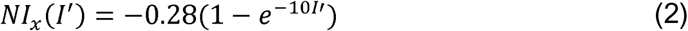

We do not assign the saturation of the *NI* with the light intensity or its gradient to the physical response of larvae since the averaged speed at which larvae move takes a fairly constant value around 4cm/min, being the small variation statistically non-significant (NS) and not obviously correlated to the intensity or the gradient of light (Fig 1E).

### Impact of directionality on phototaxis

In a natural environment, light emitted from an external cue harbors directional as well as intensity information (3). To test the relative importance of these two components, we have projected a light pattern labelled as “Tilted” (Fig 2A) in which the intensity linearly decreases along the *x*-axis, thus perpendicular to the light directionality along the *x*-axis. Contrary to the series of filters f1-f6, where both the light intensity and directionality drive larvae towards the same direction (*x*-axis), in the “Tilted” pattern, both effects are decoupled into two components: the light intensity artificially decreases in the –*y* direction because of the projected filter, while at the same time it naturally decreases along the-*x* direction as an effect of the increasing distance to the projector (S2 Fig 1). Consequently, in the “Tilted” pattern, *NI*_*y*_ mainly accounts for larval navigation due to the variation of the light intensity, while *NI*_*x*_ could be taken as a proxy of the larval navigation away from the light source. We next quantified larval navigation along the *x* – and *y* – axis independently (Fig 2B). Larval phototaxis can be explained both by the light source avoidance (Fig 2B, “Tilted” *NI*_*x*_) and by the avoidance of higher light intensities (Fig 2B, “Tilted” *NI*_*y*_). However, in the “Tilted” pattern, light directionality (*NI*_*x*_ = –0.25) has a stronger effect than intensity (*NI*_*y*_ = –0.07) in driving negative phototaxis. On the other hand, even if in the case of wildtype larvae the light-intensity-driven *NI*_*y*_ value in the “Tilted” pattern is more than three times lower than the directionality-driven *NI*_*x*_ value, it is still statistically-significantly different from the *NI*_*y*_ of the visually-blind control *glass*^*j60*^ homozygous mutant larvae, which completely lack eyes (*NI*_*y*_ *glass*^*j60*^ = –0.01, *p* – *value* = 0.015), proving that even if the wildtype *NI*_*y*_ for “Tilted” is small, it still remains light-intensity-driven taxis. Moreover, as expected, the *NI*_*x*_ for wildtype larval navigation in this “Tilted” pattern is also statistically-significantly different from the *NI*_*x*_ navigation of blind larvae (*NI*_*x*_ *glass*^*j60*^ = –0.003, *p* < 0.001). Therefore, we conclude that both the *NI*_*x*_ and the *NI*_*y*_ navigation of wildtype larvae in the “Tilted” pattern are due to the visual system and not to other effects.

**Fig 2.**
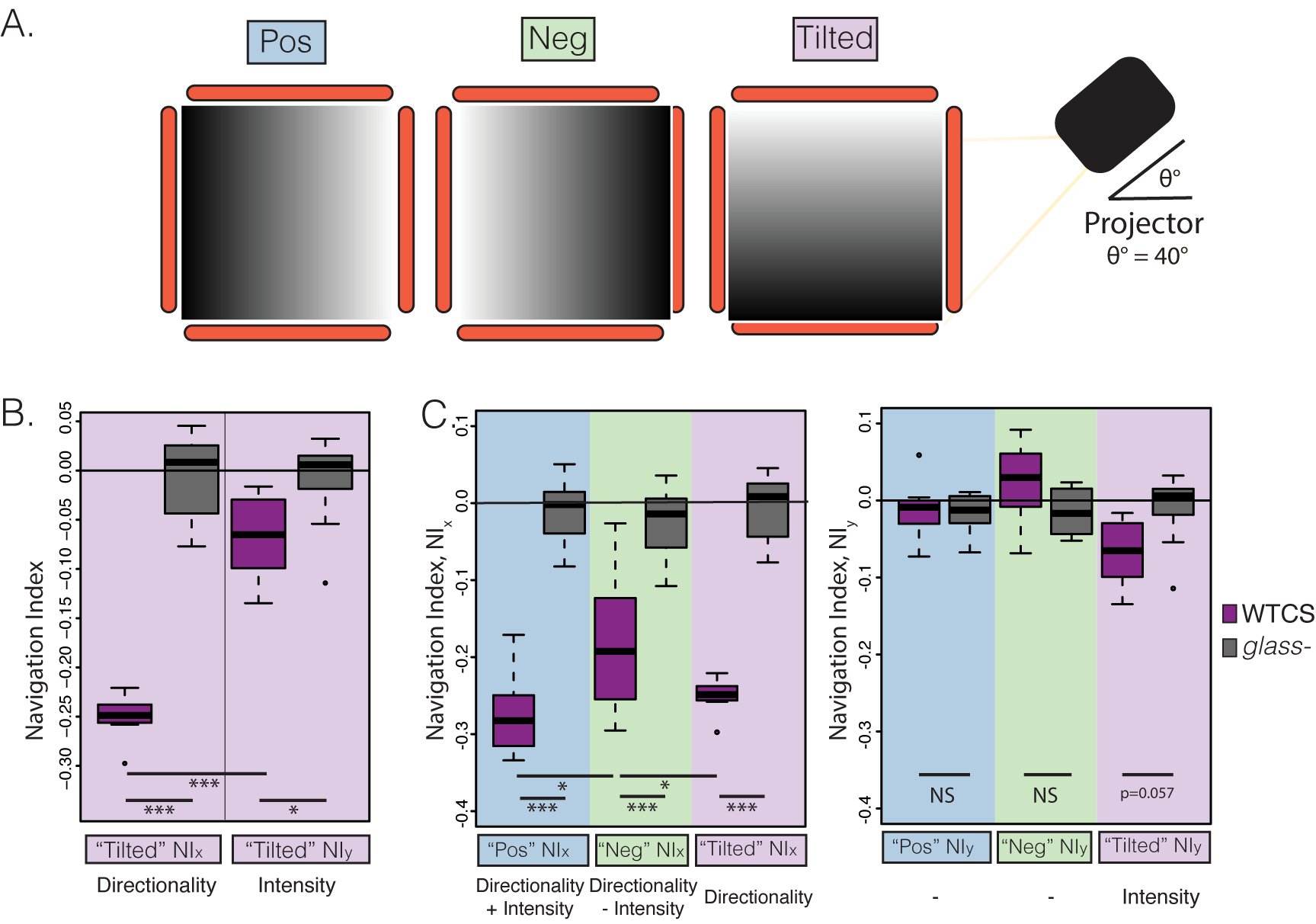
Light directionality plays a big role in larval navigation. (A) Projected filters “Pos”, “Neg” and “Tilted”, used to study the joint effect of light intensity and light directionality. All these filters were projected individually in the agarose plate and the light source was always in the +*x* side of the agarose plate as shown in Fig 1. In “Pos”, the light intensity is higher closer to the projector (right, +*x*) and decreases along the -*x* axis (increasing distance to the projector). “Neg” is the same filter but rotated 180°, where the light intensity increases as the distance from the light source increases. “Tilted” is a 90° rotation of the “Pos” filter: light intensity increases in the +*y* direction. (B) Wild-type Canton S (WTCS) Larvae in the “Tilted” filter (dark purple boxplot) have a statistically-significantly different navigation index both in the *x* direction (*NI*_*x*_ = –0.25, *p* < 0.001) and in the *y* direction (*NI*_*y*_ = –0.07, *p* = 0.0152) compared with the effectively blind *glass* mutants navigating in the same filter (grey boxplot). The directionality effect (measured in “Tilted” by *NI*_*x*_) is stronger than the light intensity effect (measured by *NI*_*y*_) as most of the larval navigation is in the *x* axis (*p* < 0.001). The were compared using the Welch t-test and corrected for multiple comparison with the Benjamini-Hochberg procedure. The range of *NIs* is given by the whiskers of the boxplots. The median (50^*th*^ percentile) is represented with the middle line and the 25^th^ and 75^th^ percentile are represented by the lower and top bands of the boxplot respectively. (C) Navigation index in the axis (*NI*_*x*_) (left graph) and in the axis (*NI*_*y*_) (right graph) for the filters showed in (A) (“Pos” in blue, “Neg” in green and “Tilted” in pink) for both WTCS (dark purple boxplots) and the blind *glass* mutant larvae (grey boxplots). Larvae presented with the “Pos” filter have the highest navigation index (*NI*_*x*_ = –0.27), as both the light intensity and light directionality drive larvae to navigate in the +*x* direction. Larvae navigating in the “Neg” filter have a lower navigation index (*NI*_*x*_ = –0.18) as both effects are driving them to navigate in opposing directions (light intensity towards +*NI*_*x*_ and light directionality towards – *NI*_*x*_). p-values are calculated with the Welch’s t-test and using the Benjamini-Hochberg procedure to correct for multiple comparison, where *p* > 0.001 is represented by ***, *p* > 0.01 ** and *p* > 0.05 by *.

To further investigate the relationship between directionality and light intensity, we have generated a pattern labelled as “Pos” (for positive) where both effects reinforce each other in the same direction and another one where they compete in opposite directions along the -*x* and +*x* axis, labeled as “Neg” (for negative) (Fig 2A). We observed that navigation is stronger for reinforcing intensity and directionality cues (“Pos”, *NI*_*x*_ = –0.27) than for competing ones (“Neg”, *NI*_*x*_ = –0.18). This also supports our finding from the “Tilted” pattern that the effect of light intensity is weaker than the directional one, since the difference in navigation indexes between “Pos” and “Neg” is just −0.09 (Fig 2C, Table 1), which we interpret as the drive towards +*x* in “Neg” (intensity) only subtracting about one third of the drive towards -*x* in “Pos” (directionality and intensity). Furthermore, the effectively blind *glass*^*j60*^ mutant larvae have a *NI* that is statistically indistinguishable from zero (Fig 2B and 2C), which proves that besides the different patterns of light, all other conditions are kept the same in all these three experiments. Table 1 provides values for experimental and simulated navigation indexes for all the projected filters.

**Table 1.**
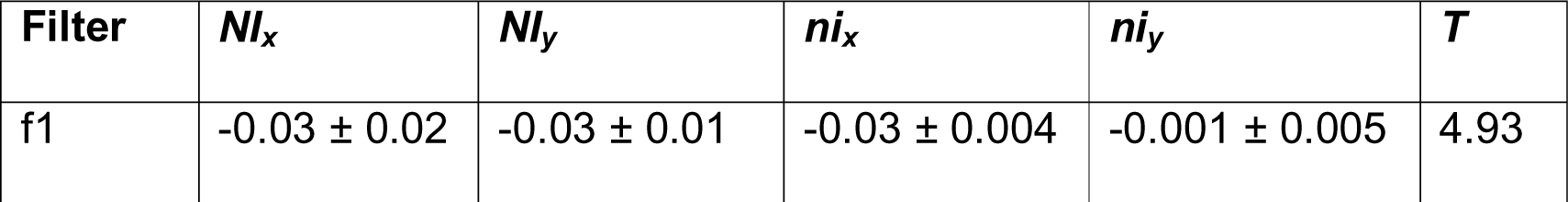
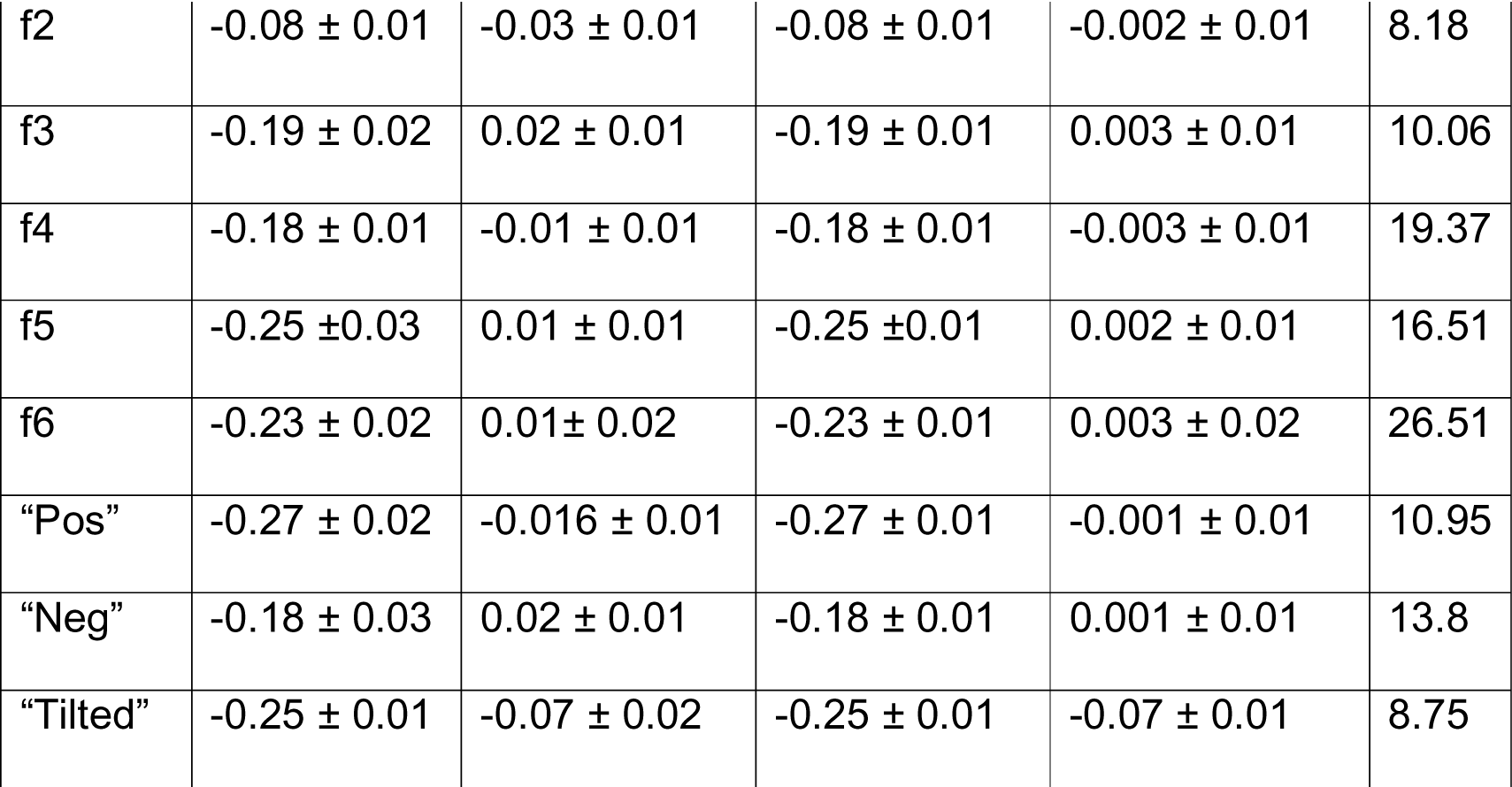
Experimental and simulated navigation indexes for all the projected patterns of lights used in the experiments (f1-f6, “Pos”, “Neg” and “Tilted”).

Experimental navigation indexes (dimensionless) in *x* ( *NI*_*x*_) and *y* ( *NI*_*y*_) directions, and the corresponding simulated navigational indexes, *ni*_*x*_ and *ni*_*y*_ for a given effective temperature *T* (W/m^2^) using 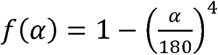. The standard deviation of the experimental *NI* was calculated for experiments for each illumination condition with around 30 larvae each. The standard deviation for the simulated *ni* was calculated with 30 simulations for each case.

### Simulation of taxis as a function of intensity and directionality

Next, we propose a mathematical model for larval phototaxis that allows us to rationalize the experimental results presented above in a unified way. Such a model must take into account two significant facts: (i) in the absence of light the observed dynamics is well described by a Markov chain resulting in a random walk characterized by *NI* = 0, and (ii) even for the higher intensities, the *NI* takes relatively low values, indicating that the external field only amounts to a small but non-negligible perturbation on the dynamics. Dealing with a perturbation has the distinctive advantage that we may assume quasie-quilibrium, same as in a diffusive regime on a physical system (24). Under these conditions, we introduce weights in the underlying Markov chain to describe the bias driven by the external field. To write an expression for these weights, we use simple biological considerations operating on the larvae; in a transition between states *r* and *r*′ in the Markov chain, we define the following weights:

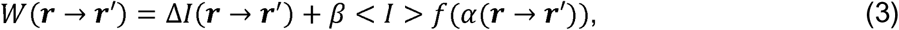

with

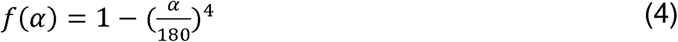

The first term of Eq (3) gives the difference in intensity experienced by larvae while taking a step from position *r* to *r*′, which depends on the gradient of the intensity, Δ*I* = *I*(*r*) – *I*(*r*′). The second term is meant to describe the directional factor. This term is proportional to the average intensity < *I* > (in units of irradiance, W/m^2^) for each projected pattern, as it deals with the increased reaction of larvae against brighter sources of light, which is hinted by the experimental results in Fig 1C. It carries an angular dependence through the function *f* of the direction *α* in the transition *r* → *r*′ (Fig 3A). We have tried different models for *f*(*α*) that will be discussed below (Materials and Methods, Determination of *f*(*α*)) and Eq (4) shows the one that yields a best fit to the experimental angular distribution probabilities. Finally, *β* is a free parameter that allows us to balance the unknown relative importance between intensity and directionality according to the experimental evidence. This parameter is obtained from a specifically-designed pattern of light (“Tilted”, Fig 2A), where the first term dominates the *NI* in one direction, while the second term dominates the *NI* in a perpendicular direction. Such a pattern provides two nearly independent experimental values for the *NI* that can be used to fit the relative importance of the two terms in Eq (3).

**Fig 3.**
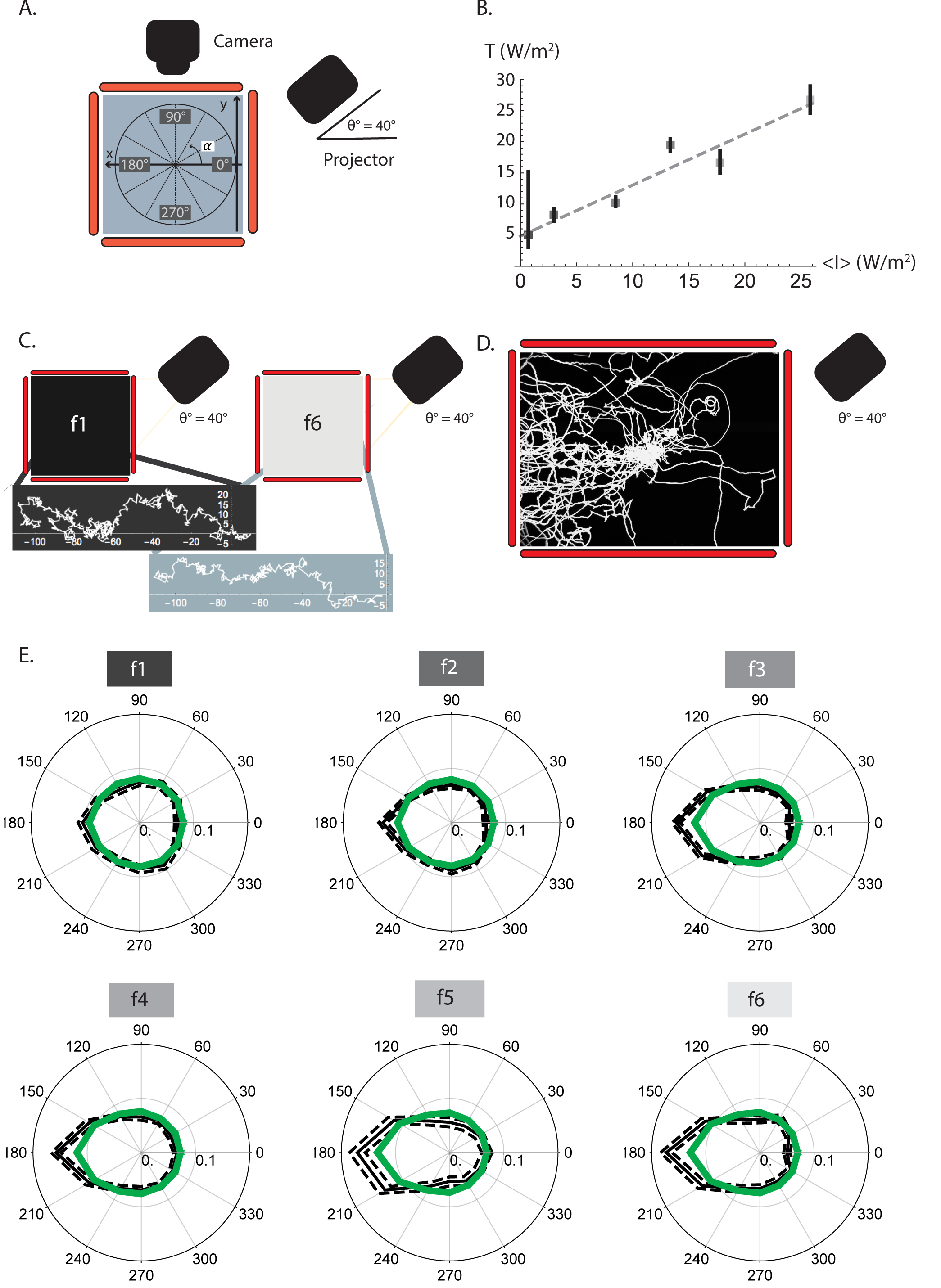
The model yields simulated larvae with similar navigation characteristics to the experimental larvae. (A). Coordinate axis used within the agarose plate to define larval movement. The angle *α* was defined within the *x* – *y* plane of the agarose plate with respect to the *x* axis, where 0° and 180° are the direction towards and away from the light source respectively. For the analysis, both the experimental and the simulated larval angular probability distributions, *P*(*α*), were binned in 30°. (B) The effective temperature *T* (W/m^2^) increases proportionally to the average intensity <*I*> (W/m^2^) of the light field. The dashed line is a linear interpolation to guide the eye. The error bars for the effective *T* have been calculated using the experimental error for the navigation index (*NI*) and calculating the *T* for *NI* plus and minus this error. (C) Simulated larval paths for f1 and f6. Each path shows one simulated larva, which starts at (0,0) and moves towards the -*x* side of the plate, avoiding the projector which is located on the +*x* side of the agarose plate (right hand side). The model yields simulated larval paths reflecting the stochastic underlying Markov chain, but also targeted navigation to get away from the light source and from the regions of high light intensities. Taxis is more targeted for higher intensity conditions (f6) same as observed in the experiments. (D) Experimental larval paths for the f6 filter (30 experimental larvae are shown); these paths are similar to the simulated ones seen in (C). (E) Comparison of the relative probability of orientation with respect to the light source (located at the right, at 0°) both for simulated (green) and experimental (grey) larvae for the different light conditions (f1, f2, f3, f4, f5 and f6). Larvae are oriented at 180° when they navigate away from the light source, and at 0° when they navigate towards the light source. The probability of orientation for both the experimental and simulated larvae has been calculated binning the possible angles in bins. The simulated larvae have been calculated using 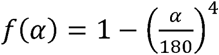. Error bars are shown for the experimental angular distributions and they were obtained from the Matlab output files from the MAGAT Analyzer.

The probability of accepting the new state *r*′ is then defined by the Boltzmann factor, *e*^‒*W*(*r* → *r*′)/*T*^ (Materials and Methods, generalized Metropolis-Hastings), where is a parameter that, following a thermodynamics simile, plays the role of an effective temperature (measured in the same units as *W*, W/m^2^). We find that the larval effective temperature has higher values for more intense light fields (f6 compared to f1, Fig 3B). We interpret the consequences of this behavior in the Discussion section. The error bar for *T* corresponding to the darkest filter (f1) is larger than for the other cases because in the absence of light, the larval movement ceases to be targeted and becomes similar to a random walk, where *T* is not meaningful anymore and cannot be determined. In mathematical terms, the effective temperature is obtained from a probability that depends on the quotient 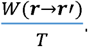 In the absence of an external field we have *W* = 0, and any value of *T* corresponds to the same probability so its value is not well defined (unbiased Markov chain).

The biased Markov chain is not time reversible (principle of detailed balance) since it is driven by an external field that imposes a definitive direction over time. However, in a similar process to the equilibration pattern followed by a thermodynamics system, we have found both in our simulations and in our experiments that important dynamical indicators, like the *NI*, reach a steady state after some initial fluctuating steps. Therefore, these indicators converge to a well-defined value and can be safely compared between simulations and experiments.

The angular part in *W*, *f*(*α*), is a dimensionless function chosen to obtain the best possible fit to the experimental angular probability distributions. We have found that a good choice for *f*(*α*) is to make it proportional to a power of the angle *α*^*n*^, where *α* is the angle formed in the plane of the agarose plate between the attempted direction *r*′ and the *x* axis (Fig 3A), and the power *n* can be taken as a free parameter that is chosen to obtain the best fit to experiments. In particular, we have found that *n* = 4 is an optimal value. Possible choices for *f*(*α*) are described in more detail below (Materials and Methods, Determination of*f*(*α*)).

Our mathematical model yields more targeted paths when the intensity and its gradient are higher (Fig 3C, f6 compared to f1) which leads to higher values for *NI*. Furthermore, simulated larval paths look similar to the experimental ones (Fig 3D), which indicates that not only averaged values like the *NI* can be successfully compared between simulations and experiments. In fact, the similarity of the angular distribution for the experimental larvae and the simulated ones proves that our model quantitatively reproduces experimental results under different light conditions; Fig 3E shows the agreement between experimental (grey curves) and simulated (green curves) angular distributions obtained with the non-linear 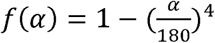 for filters f1 to f6.

## Discussion

### Larvae respond differently to different intensities and light gradients

Our results show that larval navigation depends on light intensity and its gradient and that larvae navigate more efficiently (larger *NI*) when the light intensity is higher and when the gradient of the intensity is steeper. However, this non-linear behavior (Eqs (1) and (2)) saturates for intensities higher than *I* > 20 W/m^2^ (Fig 1C) and for intensity gradients higher than *I*′ > 0.2 W/m^2^/cm (Fig 1D). As commented above, we do not assign such a saturation behavior to a lack of physical response from the larvae since the mean velocity is nearly independent of the illumination conditions, but rather to a limited capacity for processing information in the underlying neural network in the larval brain. From a biological point of view, larvae make decisions to take a step from position *r* to *r*′ or not, based on the *local* conditions surrounding them. These conditions are the only ones that larvae can probe with their limited visual organ and the only ones that they can process in their brains without requiring an expensive memory process to record a string of magnitudes all along their paths. Saturation regarding the light intensity can be understood from a limited ability to process the input signal of too many photons. On the one hand, the experimental evidence that larvae navigate differently depending on the gradient of the intensity implies that larvae must read the gradient of the field of light (*I*′). Such an operation needs to measure the intensity in two close points and then proceed to compare them. Therefore, it involves a *memory* process if it is to be done at two subsequent times along the larval path.

Similarly, in the model, whether a transition in the Markov chain happens or not is based on variables that can be locally obtained using only the larval starting position.

### Distinct larval directional tendency to move away from the light source

The combination of our experiments with our mathematical model shows the detailed relationship between light intensity and directionality for larval phototaxis for the first time. The analysis of the “Tilted” pattern by our mathematical model tells us that the main reason for larval phototaxis, in about a one to three ratio, is to get away from the source of light, rather than to simply move to darker regions. This is in part due to the saturation process shown in Fig 1C and 1D, that limits the effect of the first part in Eq (3). The weights defined in our model show, in a quantitative way, that larvae can process the relative orientation of light. Such a capacity to discriminate different angles explains the larval ability to move maximizing the distance to the source of light by simply giving a higher probability to angles around 180° (away from the light source) than to angles around 0° (towards the light source).

### Larval phototaxis and the proposed mathematical model

An interesting feature of our mathematical model is that it only requires three adjustable parameters: the relative balance between intensity and directionality *β*, Eq (3); the power *n* telling how sensitive the larvae is to changes in the light direction, Eq (4); and the effective temperature *T* that determines the stochastic exploration (see Eq 5 below). The reason for needing so few parameters is probably linked to the general principles governing the generalized Metropolis-Hastings algorithm, which takes care of the statistical behavior in a way that is known to work well for many different complex systems found in nature. The model is based in principles so well accepted in different contexts that except for particular details in Eq (3), it should work for other organisms and other sensory cues.

The related Metropolis-Hastings algorithm has been successfully used to efficiently locate *global* minima of combinatorially-complex objective functions such as the travelling salesman problem (17). In contrast, we remark that our biological experiments mostly bring information about larval decisions taking into account *local* data (intensities and gradients in the immediate surroundings of the organism) and proceed with a limited amount of neural circuitry. Therefore, the weights governing the simulation in Eq (3) should be considered as a local solution to the problem rather than a global one.

The value of the effective *T* in the simulations is adjusted so that the currents of larvae going towards the light source and in the opposite direction match the experimental *NI* measured under some particular light conditions. Therefore, this *T* determines a quasi-equilibrium condition, similar to the one found in a chemical reaction where reactants convert into products, and vice versa, in ratios that match the actual production at a given temperature. On the other hand, such a parameter lends itself to a biological interpretation; it controls the larval probability of taking risks by either going to higher intensity regions or by getting closer to the source of light. Such a behavior is known to be a useful way to avoid being trapped in local minima, as it has been proved when simulated annealing has been applied to find the global minimum of a given objective function. Our results show that the effective temperature grows with light intensity and its gradient. These are the conditions that from a biological point of view should require more vigorous action from the organism to quickly find a more convenient position. In turn, when conditions are not so harsh, organisms prefer taking conservative decisions, hardly moving to worse regions in order to explore their environment more efficiently.

Regarding the different models that we have tried for *f*(*α*), the functions that reproduce the experimental data better are the power-like ones. Linear functions of the angle do not agree well with experiments; all reasonable candidates have been highly non-linear functions. Therefore, the larval behavior reveals a complex and rich neural network behind the process of taking decisions, which works on a non-linear function, which is a common feature to neural circuits organized in layers (25).

So far, the model does not take into account the larval dimensions. However, it would be possible to add terms to the weights *W*(*r* → *r*′) in Eq (3) to take into account the size of the larva, for example, by artificially increasing the value of these weights when two larvae would overlap on the new position *r*′. In this case, we could take into account the fact that they cannot go to places already occupied by other larvae and even the attraction or repulsion between individuals could be modeled (26, 27). This would open the possibility to study simulated group behavior, although at a higher computational cost.

An additional feature of larval taxis, studied for chemotaxis, is *weathervaning*, which is defined as miniature head-sweeping during runs resulting in curved tracks (28). Whether *weathervaning* plays a role in phototaxis or not remains unclear. Since our mathematical model is based on the local values of weights for the underlying Markov chain, *weathervaning* is not directly taken into account. The model is based on larval runs and turns with a single underlying mechanism governed by the weights in Eq (3). Moreover, (29) have studied *weathervaning* for larval chemotaxis to conclude that it is the least crucial navigational parameter according to their model.

### Relevance for the neuronal network involved in larval phototaxis

Mathematical models of larval navigation provide the first step towards understanding the underlying mechanisms that operate in the larval neuronal network to lead to decision-making. Next steps in understanding the neuronal basis of visual navigation may include to combine current information of the connectome with behavioral data and to correspondingly adapt a mathematical model (30, 31). This generalized Metropolis-Hastings-based model could also be used for other stimuli, the only requirement being that the intensities and gradients of these stimuli should be measured and quantified properly to define the details of the weights in Eq (3). Once these weights have been found for other stimuli, multi-sensory experiments could be carried out to see if these factors are additive towards larval navigation as suggested in (12).

## Materials and Methods

### Fly strains

Wild-type Canton S (WTCS) *D. melanogaster* larvae (courtesy of R. Stocker), and *glass*^*60j*^ mutants (Bloomington 509) were used for these experiments. All the fly stocks were kept at 25°C in a 12-hour light-dark cycle. The stocks were fed with a conventional cornmeal medium containing molasses, fructose and yeast.

### Behavioral experiments

Larvae were selected for experiments after four days since the egg-laying of the parental flies, ensuring that they would correspond to the 3^rd^ larval stage (L3). Larvae were kept for at least 10 minutes in the dark with food before the phototaxis experiments were carried out. Thirty larvae were isolated from the food for each experiment and placed in water droplets with a paintbrush. The maximum time for the larval selection was 10 minutes and it was done under red light conditions. The larvae were left in the agarose plate without food and their tracks were recorded for 11 minutes. The first minute was not taken into account to let the larvae get used to the new conditions. The behavior experiments were always carried out within the larval 12 light-hours.

### Tracking system

The experimental setup consists of a 23×23 cm agarose plate where the larvae can move freely (Fig 1A). Larval movements were recorded with a Basler acA2500-14gm camera equipped with a 1:14/12.5 mm Fujinon lens and placed directly above the tracking arena. The lens was incorporated with a red filter (635 nm, Qualimatest SA, Geneva, Switzerland). The agarose plate was illuminated with red LEDs that do not influence larval behavior but enable the image recollection with the camera (Fig 1A).

An EB U04 projector was located at *x* = 36.5 cm,*y* = 0 cm, *z* = 25 cm. The projector was equipped with a 2" Square BG40 coloured glass bandpass filter 335 – 610 nm and was placed forming a 40° angle with respect to the *x* – *y* plane formed by the agarose plate (zenithal angle, *θ*, Fig 1A).

The custom-made LabView software (32,3) was used to record the larval movies.

### Light intensity measurement on the agarose plate

Different light patterns were projected to obtain the different experimental scenarios against which the simulations could be validated. The intensity field varied in a different way in all of them. f1-f6 were uniform filters where the intensity variation was merely due to variation of the photon flux with the distance to the projector (Fig 1B and S1 Fig 1).In the patterns “Pos”, “Neg”, and “Tilted” used to explore directionality, an artificial modulation in the light gradient along the *x* or *y* directions was introduced and therefore there was a steeper change in light intensity. In “Pos”, the maximum light intensity was closer to the light source, same as in all the f1-f6 patterns, but the light intensity decreased along the axis with a gradient that was about 5 times steeper (S1 Table 1 and S2 Table 1). “Neg” was a 180° rotation of “Pos”; therefore, the intensity field decreased along the axis and the brightest area was located further away from the light source. “Tilted” was a 90° rotated version of the “Pos” pattern. In this case, the light gradient artificially varied along the y direction, therefore the intensity field decreased along the –*y* axis (Fig 2A, S2 Fig 1C).

Light intensities created by the different projected patterns were measured on the agarose plate using an *Ocean Optics USB400* spectrometer. Three equally-spaced points along the *x* axis (*y* = 0) were measured for the filters used to study light intensity (f1-f6, S1 Fig 1) and nine points equally covering the *x* and *y* directions for the patterns related with directionality (“Pos”, “Neg”, and “Tilted”, S2 Fig 1). Several measurements were taken for each point on different days and standard errors were calculated. The total intensity (W/m^2^) was obtained by integrating these spectra between 380 and 570 nm to include the blue and green wavelength regions of the spectrum relevant for Rh5 and Rh6 absorption spectra, but to exclude the red one (S1 Fig 1 and S2 Fig 1). Integrals have been performed by first defining an interpolating polynomial going through all the experimental points, and then using an accurate Gaussian-Konrod rule for integration (33).

The expected variation of light intensity on the plate by a uniform source of light is described by assuming a steady rate of generation of photons. For the actual parameters of the geometrical setup, this has the implication of an approximate linear variation, which has been corroborated by measuring intensities on the plate (S1 Fig 1 and S2 Fig 1). Therefore, the spatial variation of intensities has been represented by a linear fit with

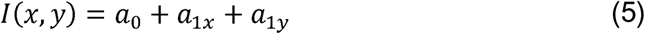

where *a*_0_ is the value at *x* = *y* = 0 and *a*_1*x*_, *a*_1*y*_ are the slopes along the *x* and *y* directions. Values for these coefficients are given in S1 Table 1 and S2 Table 1.

The air conditioning was turned on at 25°C during the experiments to ensure a constant temperature in the agarose plate. Measurements of the temperature on the plate always yielded temperatures in the interval between 25°C and 26°C.

### Tracking data analysis

The acquired images of the larval tracks were analyzed with the MAGAT Analyzer (3). The features of each larva (head, tail and midline) were extracted from the videos and these data were analyzed using a custom-made software written in MATLAB (34).

Statistical analysis of the data was calculated using the Welch's unpaired t-test to compare results with different genotypes and a regular unpaired t-test was used to compare larvae with the same genotype. The Benjamini-Hochberg procedure was applied to correct for multiple comparison. The statistical difference of results compared with zero was calculated using a one-sample t-test.

### Generalized Metropolis-Hastings chains

In our simulations, we assign transition probabilities between states in the Markov chain according to the Boltzmann distribution. Probabilities are assigned in the following way ( 35):

1. A description of possible system configurations and the options presented the system. These are determined by giving:

a. The initial position of the larvae, *r* = (*x*, *y*) and the final attempted position, *r*′ = (*x*′, *y*′), *where x*′ = *x* + Δ*x and y*′ = *y* + Δ*y*
b. The light intensity at both points: *I*(*r*) *and I*(*r*′)
2. A generator of random changes in the configurations. We chose the next position using two independent Gaussian deviates with zero mean (Δ̄ = 0) and standard deviation one (*σ* = 1) for the independent increments in the *x* and the *y* direction, Δ*x* and Δ*y* respectively. This defines a discretetime continuous-space Markov chain of transition kernel *k*_0_ (*r*, *r*′) = 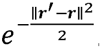. The standard deviation sets up a length scale that we adjust to the observation that the larvae approximately advance a distance equivalent to the length of its body in about ten moves. Therefore, the standard deviation is equivalent to approximately 0.1 mm.
3. *The larval local moves are described by a discrete-time continuous-space Markov chain of transition Kernel:*

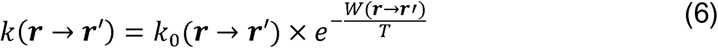

where *W*(*r* → *r*′) = Δ*I*(*r* → *r*′) + *w* < *I* > *f*(*α*(*r* → *r*′)). Algorithmically, the new *r*′ is chosen as follows:

- Choose *r*′ according to the Gaussian model (*k*_0_), *r*′ = *r* + *μ*, where *μ* is a bivariate Gaussian deviate with zero mean (Δ̄ = 0) and standard deviation in both dimensions
- If *W*(*r* → *r*′) < 0, then *r*′ is accepted with probability *P* = 1
- If *W*(*r* → *r*′) > 0, then *r*′ is accepted with probability 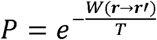
4. *A control parameter T*. This parameter controls the weights so that the simulation can reproduce the experimental navigation index for a given light intensity pattern. *T* has irradiance units, same as *W*, and in a thermodynamics system it would be the equilibrium temperature. High values of *T* leads to small values of the exponential weight 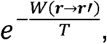, and the related transition step will occur with a probability near to 1 for positive *W*.

The experimental recording stops tracking larvae that hit the border of the agarose plate. Accordingly, we introduce a similar boundary condition in our simulations and we stop tracking larvae that after *N* accepted steps have reached a distance to the origin greater or equal to 1150 *σ* ≈ 11.5 cm. Depending on the illumination conditions, this usually happens after a few thousands accepted steps. At that point, the Markov chain has an absorbing state. However, we checked that the averaged *NI* and the angular probability distributions reached a quasi-stationary state before the experiment is terminated. For a given light pattern, each simulation was carried out with 30 individual larvae, and the average and its standard error have also been computed for a 30 larvae ensemble, making sure that the given *NI* is statistically significant.

### Determination of *f*(*α*)

The angular part in equation (3) has been modelled to take into account the experimental angular distributions. The angle *α* is measured with respect to the *x* axis, being 0° the direction towards the projector and 180° the direction away from it (Fig 3A). Two types of models for *f*(*α*) were tried: power-like models proportional to *α*^*n*^ and models based on *cos*^*n*^, taking into account that cos(*α*) = Δ*x*/Δ*l* (S3 Fig 1). Each model was assessed calculating the standard deviation of the angular probability distribution of the simulated paths compared to the experimental ones. Both the experimental and simulated paths were binned in 30° angles and the probabilities for both experimental and simulated cases were compared. The best fit to experiments across all the different projected patterns was found for *f*(*α*) = 1–(*α*/180)^4^, as shown in S3 Table 1,where we give the root-mean-squared (RMS) deviation between experimental and simulated angular distributions for all models tried for *f*(*α*).

### Determination of *β*

The value for the parameter *β*(*β* = 1.4/100) was determined taking the “Tilted” pattern as a case where the two terms of the objective function (intensity and directionality) are most decoupled. Consequently, the *NI*_*y*_ in the “Tilted” pattern was simulated assuming that only the first term in the objective function would exist. That procedure yields a value for the effective *T*. Afterwards, the value for *NI*_*x*_ was used to find a value for the parameter *β*. As a final consistency check, both the *NI*_*y*_and the *NI*_*x*_ were simultaneously recalculated using the two parts of the objective function obtaining a refined value for that fits the two available experimental values at the same time.

## Acknowledgements

LdA would like to thank Tim-Henning Humberg for help with the experimental set up and fruitful discussions and comments, as well as other members of Prof. Sprecher’s lab for their support with the experiments. We thank members of Prof. Mazza’s and Prof. Senn’s groups, in particular Martin Wiechert for useful comments on setting up the mathematical model. This work was supported by the SystemsX.ch initiative.

## Supporting information

**S1 Fig 1. Light spectrum along the agarose plate for the f1 – f6 filters.** (A-F) The light intensity was measured with an OceanOptics USB400 spectrophotometer in the agarose plate in three points for the f1 – f6 filters. Light intensities were measured for all the filters along the *x* axis of the agarose plate at three points: (1) one closer to the projector, at (11.5,0) cm (white square), (2) in the center of the agarose plate (0,0) cm (gray square) and (3) in the point furthest away from the projector (–11.5,0) cm (black square). Replicates of each of the measurements were taken on different days. (A’-F’) Light intensity (*I*) is given in *μ*W/cm^2^/nm as a function of the wavelength *λ* in nm. The light intensity measured on these three points was plotted and integrated between 380 nm and 570 nm (vertical lines), which is the biologically relevant wavelength range for the larvae. (A”-F”) The variation of light intensity along the *x* axis of the agarose plate was modelled with a linear regression for each of the filters.

**S1 Table 1. Light measurements for the f1 – f6 projected filters.** The three measured points along the *x* axis of the agarose plate for filters f1 – f6 (S1 Fig1 A-F) were used to model the variation with a linear regression. Light intensity is quantified with a least-square polynomial fit to intensities. The slope of the linear variation of intensities (*a*_1*x*_*x* in W/m^2^/cm) and the intercept (*a*_0_ in W/m^2^) can be seen in this table for each of the filters.

**S2 Fig 1. Light spectrum along the agarose plate for the directionality patterns.** (A-C) The light intensity was measured as in S1 Fig 1 for the “Pos”, “Neg” and “Tilted” patterns in nine points forming a homogenous grid in *x* and *y*. The gradients of light intensities along the axis *x* and along the *y* axis were obtained from that grid. Replicates of each measurement were taken on different days. (A’-C’) Same as in S1 Fig 1 for “Pos”, “Neg”, and “Tilted”. (A”-C”) Variation of light intensity along the *x* axis on the agarose plate for “Pos” and “Neg” and along the *y* axis for “Tilted”. A linear regression for each of the patterns was found to describe well the pattern of intensities. (A”’-C”’) Contour plot for each of the projected filters.

**S2 Table 1. Light measurements for the projected directionality patterns.** Light intensity is quantified with a least-square polynomial fit to intensities, *a*_0_ + *a*_1*x*_*x* + *a*_1*y*_*y* This table shows the values for the linear regression parameters: the slope of the linear variation of the light intensity along the *x* axis (*a*_1*x*_ in W/m^2^/cm), along the *y* axis (*a*_1*y*_ in W/m^2^/cm) and the intercept (*a*_0_ in W/m^2^) for each of the projected patterns.

**S3 Table 1. Comparison of the different models for f(β).** Simulations were carried out using different models for *f*(*β*) and tested against the f1 – f6 filters. Simulations for each model were carried out 30 times and the experimental ones were calculated doing 10 experiments with around 30 larvae each. Both the experimental and simulated angular probability distributions were binned in 30° angles. Each model was assessed by calculating the root mean squared deviation (RMS) between the experimental and simulated angular probability distributions for the different binned angles from 0° to 180°. Then, the average of the RMS for all the filters was compared for each model of *f*(*α*). The smallest overall RMS was obtained with *f*(*α*) ∝ 1 – *α*^4^.

**S3 Fig 1. Geometrical diagram for the directionality part of the cost function.** The source of light is approximated by a plane, F. The light came from right (+*x*) to left (–*x*). The angular distribution is a function of the angle of the direction of the larva, *α*, which is a function of the displacement *α* = arctan 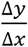. The increment in the distance to the plane F depends on the displacement Δ*x* = Δ*l cosα*, where 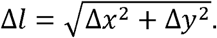.

## References

1. Berg HC, Brown DA. Chemotaxis in Escherichia coli analysed by threedimensional tracking. Nature. 1972;239(5374):500–4.

2. Lockery SR. The computational worm: spatial orientation and its neuronal basis in C. elegans. Curr Opin Neurobiol. 2011;21(5):782–90.

3. Kane EA, Gershow M, Afonso B, Larderet I, Klein M, Carter AR, et al. Sensorimotor structure of Drosophila larva phototaxis. Proc Natl Acad Sci USA. 2013;110(40):E3868–77.

4. Humberg TH, Sprecher SG. Age‐ and Wavelength-Dependency of Drosophila Larval Phototaxis and Behavioral Responses to Natural Lighting Conditions. Front Behav Neurosci. 2017;11.

5. Louis M, Huber T, Benton R, Sakmar TP, Vosshall LB. Bilateral olfactory sensory input enhances chemotaxis behavior. Nat Neurosci. 2008;11(2):187–99.

6. Gomez-Marin A, Stephens GJ, Louis M. Active sampling and decision making in Drosophila chemotaxis. Nat Commun. 2011;2:441.

7. Ohashi S, Morimoto T, Suzuki Y, Miyakawa H, Aonishi T. A novel behavioral strategy, continuous biased running, during chemotaxis in Drosophila larvae. Neurosci Lett. 2014;570:10–5.

8. Schulze A, Gomez-Marin A, Rajendran VG, Lott G, Musy M, Ahammad P, et al. Dynamical feature extraction at the sensory periphery guides chemotaxis. Elife. 2015;4.

9. Luo L, Gershow M, Rosenzweig M, Kang K, Fang-Yen C, Garrity PA, et al. Navigational decision making in Drosophila thermotaxis. J Neurosci. 2010;30(12):4261–72.

10. Lahiri S, Shen K, Klein M, Tang A, Kane E, Gershow M, et al. Two alternating motor programs drive navigation in Drosophila larva. PLoS One. 2011;6(8):e23180.

11. Klein M, Afonso B, Vonner AJ, Hernandez-Nunez L, Berck M, Tabone CJ, et al. Sensory determinants of behavioral dynamics in Drosophila thermotaxis. Proc Natl Acad Sci USA. 2015;112(2):E220–9.

12. Gepner R, Mihovilovic Skanata M, Bernat NM, Kaplow M, Gershow M. Computations underlying Drosophila photo-taxis, odor-taxis, and multi-sensory integration. Elife. 2015;4.

13. Hernandez-Nunez L, Belina J, Klein M, Si G, Claus L, Carlson JR, et al. Reverse-correlation analysis of navigation dynamics in Drosophila larva using optogenetics. Elife. 2015;4.

14. Gunther MN, Nettesheim G, Shubeita GT. Quantifying and predicting Drosophila larvae crawling phenotypes. Sci Rep. 2016;6:27972.

15. Mazza C, Benaïm M. Stochastic dynamics for systems biology. Boca Raton FL: CRC Press, Taylor & Francis Group; 2014. xii, 260 p. p.

16. Klein M, Krivov SV, Ferrer AJ, Luo L, Samuel AD, Karplus M. Exploratory search during directed navigation in C. welegans and Drosophila larva. Elife. 2017;6.

17. Kirkpatrick S, Gelatt CD, Jr., Vecchi MP. Optimization by simulated annealing. Science. 1983;220(4598):671–80.

18. Wystrach A, Lagogiannis K, Webb B. Continuous lateral oscillations as a core mechanism for taxis in Drosophila larvae. Elife. 2016;5.

19. Mazzoni EO, Desplan C, Blau J. Circadian pacemaker neurons transmit and modulate visual information to control a rapid behavioral response. Neuron. 2005;45(2):293–300.

20. Keene AC, Mazzoni EO, Zhen J, Younger MA, Yamaguchi S, Blau J, et al. Distinct visual pathways mediate Drosophila larval light avoidance and circadian clock entrainment. J Neurosci. 2011;31(17):6527–34.

21. Green P, Hartenstein AY, Hartenstein V. The embryonic development of the Drosophila visual system. Cell Tissue Res. 1993;273(3):583–98.

22. Sprecher SG, Desplan C. Switch of rhodopsin expression in terminally differentiated Drosophila sensory neurons. Nature. 2008;454(7203):533–7.

23. Sprecher SG, Pichaud F, Desplan C. Adult and larval photoreceptors use different mechanisms to specify the same Rhodopsin fates. Genes Dev. 2007;21(17):2182–95.

24. Feynman RP, Leighton RB, Sands ML. The Feynman lectures on physics. Reading, Mass.; London: Addison-Wesley Pub. Co.; 1963. 3 volumes p.

25. Rutishauser U, Douglas RJ, Slotine JJ. Collective stability of networks of winner-take-all circuits. Neural Comput. 2011;23(3):735–73.

26. Otto N, Risse B, Berh D, Bittern J, Jiang X, Klambt C. Interactions among Drosophila larvae before and during collision. Sci Rep. 2016;6:31564.

27. Niewalda T, Jeske I, Michels B, Gerber B. 'Peer pressure' in larval Drosophila? Biol Open. 2014;3(7):575–82.

28. Gomez-Marin A, Louis M. Multilevel control of run orientation in Drosophila larval chemotaxis. Front Behav Neurosci. 2014;8:38.

29. Davies A, Louis M, Webb B. A Model of Drosophila Larva Chemotaxis. PLoS Comput Biol. 2015;11(11):el004606.

30. Larderet I, Fritsch PMJ, Gendre N, Neagu-Maier GL, Fetter RD, Schneider-Mizell CM, et al. Organization of the Drosophila larval visual circuit. Elife. 2017;6.

31. Sprecher SG, Cardona A, Hartenstein V. The Drosophila larval visual system: high-resolution analysis of a simple visual neuropil. Dev Biol. 2011;358(1):33–43.

32. Gershow M, Berck M, Mathew D, Luo L, Kane EA, Carlson JR, et al. Controlling airborne cues to study small animal navigation. Nat Methods. 2012;9(3):290–6.

33. Wolfram. Mathematica [Available from: https://www.wolfram.com/mathematica/.

34. Mathworks. Matlab 2017 [31st October 2017]. Available from: https://ch.mathworks.com/products/matlab.html.

35. Press WH, Teukolsky SA. Numerical recipes: Does this paradigm have a future? Comput Phys. 1997;11(5):416–24.

